# Cortical Response to Acute Implantation of the Utah Optrode Array in Macaque Cortex

**DOI:** 10.1101/2025.01.13.632843

**Authors:** Adrián Villamarin-Ortiz, Christopher F. Reiche, Frederick Federer, Andrew M. Clark, John D. Rolston, Cristina Soto-Sánchez, Eduardo Fernandez, Steve Blair, Alessandra Angelucci

**Author notes:** E. Fernandez, S. Blair and A. Angelucci contributed equally to this work and share senior authorship.

## Abstract

Optogenetics has transformed the study of neural circuit function, but limitations in its application to species with large brains, such as non-human primates (NHPs), remain. A major challenge in NHP optogenetics is delivering light to sufficiently large volumes of deep neural tissue with high spatiotemporal precision, without simultaneously affecting superficial tissue. To overcome these limitations, we recently developed and tested *in vivo* in NHP cortex, the Utah Optrode Array (UOA). This is a 10×10 array of penetrating glass shanks, tiling a 4×4mm^2^ area, bonded to interleaved needle-aligned and interstitial µLED arrays, which allows for independent photostimulation of deep and superficial brain tissue. Here, we investigate the acute biological response to UOA implantation in NHP cortex, with the goal of optimizing device design for reduced insertion trauma and subsequent chronic response. To this goal, we systematically vary UOA shank diameter, surface texture, tip geometry, and insertion pressure, and assess their effects on astrocytes, microglia, and neuronal viability, following acute implantation. We find that UOAs with shanks of smaller diameter, smooth surface texture and round tips cause the least damage. Higher insertion pressures have limited effects on the inflammatory response, but lead to greater tissue compression. Our results highlight the importance of balancing shank diameter, tip geometry, and insertion pressure in UOA design for preserving tissue integrity and improving long-term UOA performance and biocompatibility.

## 1. Introduction

Optogenetics has transformed the study of neural circuit function^[1]^, and offers great potential for clinical applications^[2]^. Unlike conventional electrical microstimulation^[3]^, optogenetics allows for manipulations of neural activity in a cell-type specific manner at physiologically relevant time scales. While progress in the application of cell type-specific optogenetics to species with large brain sizes, such as non-human primates (NHPs), has lagged behind that in the mouse^[4]^, recent advances in viral technology^[5–10]^ are rapidly opening up new opportunities to study neural circuits in NHPs and even humans. Extending optogenetics to NHPs is important for understanding neural circuit function and dysfunction in human neurological and psychiatric disorders^[11–12]^, as this species is the closest to humans and provides an essential technology testbed for the development of optogenetic therapies^[13–16]^.

Despite recent advances, a primary limitation in the application of optogenetics to NHPs has been the difficulty of delivering light of sufficient irradiance to deep neural tissue across large brain volumes, to modulate relevant circuits and behavior. Photostimulation through the brain surface, a widely used light delivery approach^[17–20]^, limits photoactivation to a depth of <1mm, due to light scattering and absorption within brain tissue^[21]^. Moreover, this approach does not allow to restrict photostimulation to deep neural tissue without also activating superficial tissue. Penetrating probes, instead, allow for focal light delivery at depths >1mm^[22–23]^, but current devices applicable to NHP studies are limited to a single optical fiber^[24–26]^, or few (up to 4) optical fibers as in the commercially available Plexon opto-probe^[27]^, which only allow for a total activation volume of a few hundred microns in diameter.

To overcome the limitations of available devices for optogenetic stimulation in NHPs, we recently developed, and tested *in vivo* in NHP cortex, the Utah Optrode Array (UOA)^[28–29]^. Inspired by the widely used, FDA-approved, Utah Electrode Array (UEA) for microcurrent delivery^[30]^, the UOA is capable of independently delivering light to deep and superficial brain tissue over a large volume with high spatio-temporal precision. It consists of a 10×10 array of penetrating glass needle shanks, acting as light waveguides to deep tissue, tiling a 4×4mm^2^ area, bonded to an electrically-addressable µLED array independently delivering light through each shank. A second 9×9 µLED array is interleaved with the needle-aligned µLED array and matrix-addressed for independent surface stimulation^^[28, 31–32]^^. *In vivo* testing in acute anesthetized NHP visual cortex demonstrated the UOA allows for spatiotemporally patterned photostimulation of deep cortical layers with sub-millimeter resolution over a large volume, and that this selectivity can be scaled up by varying the number of simultaneously activated µLEDs and/or the light irradiance^[29]^.

Unfortunately, intracortical probes face significant biocompatibility challenges, as implantation of these devices disrupts the blood-brain barrier, initiating an inflammatory response. This response often leads to the formation of a glial scar surrounding the device shanks, which can impair their long-term functionality. Additionally, the extent of this reaction may escalate to a higher number of shanks in the tissue, further compromising the probe functionality over time^^[33–34]^^. While some studies have investigated flexible microelectrodes as an alternative to reduce inflammatory responses by conforming more closely to natural brain tissue movements^[35–38]^, these designs lack the necessary rigidity and may bend during implantation, compromising the precision of targeting the intended sites^[39]^. In contrast, the UOA, with its structural robustness and established use in intracortical applications, may remain the preferred choice for targeted photostimulation, particularly in large brains.

Understanding and enhancing the biocompatibility of the UOA is essential to mitigate adverse responses and improve long-term probe performance. Acute tissue responses are particularly valuable, as they offer early insights into initial tissue reactions upon device implantation. By minimizing insertion trauma during this phase, the risk of subsequent chronic responses may be significantly reduced^[40]^.

This study aimed to elucidate the acute biological response to UOA implantation in a NHP model. We systematically varied key physical and mechanical parameters of the UOA, including shank diameter, surface texture, tip geometry, and insertion pressure, to assess their effects on astrocytes, microglia, and neuronal preservation following acute implantation. Investigating how these design features influence acute responses will guide a more comprehensive optimization of intracortical device design and insertion procedures, potentially minimizing tissue damage while ensuring sustained functionality over extended periods.

## 2. Results

### 2.1. The Utah Optrode Array (UOA)

The UOA is a 10 x 10 array of penetrating glass optical needle shanks with customizable length (up to 2.5 mm) and width (60-120 µm) on a 400 µm pitch, tiling a 4×4 mm^2^ area (**Fig. 1A-B**). The penetrating shanks deliver light to deep brain tissue. In its “active” form, the device is bonded to an electrically addressable μLED array which allows for independent light delivery through each shank through the shank tip^[31–32]^, and to an interleaved 9×9 µLED array for surface illumination between the shanks. The UOA is based on the geometry of the UEA^[30]^, but its shank width is smaller than that of the UEA at the base (150µm), and, unlike the UEA, does not taper (to prevent light leakage), therefore at the tip it is slightly wider than the UEA. Its tip geometry can be controlled through the fabrication process, from sharp (but still wider than the UEA) to round (**Fig.1C**). Tip geometry affects both the light coupling efficiency and the light emission profile, with more rounded tips giving higher peak irradiance and deep illumination, and sharper tips giving less irradiance and a lateral illumination profile^[28]^. However, sharper tips require less insertion force than round tips to penetrate brain tissue, with consequent reduced tissue trauma and vascular damage^[41]^. To reduce the surface roughness of the shanks resulting from prior fabrication steps the UOA must undergo a high temperature annealing/reflow step^[31–32]^, and the longer the reflow phase is performed, the smoother the shanks become. Smoother surfaces reduce light scattering in the shank. However, the annealing step also causes a rounding of the corners at the shank tip, which therefore becomes more rounded (hemispherical) in shape, the smoother the shank surface becomes. More rounded tips penetrate tissue less easily than sharper tips, presumably causing more damage. Therefore, the fabrication process needs to achieve a fine balance between minimizing surface roughness, for more efficient light coupling, and tip roundness, for easier tissue penetration.

**Figure 1.**
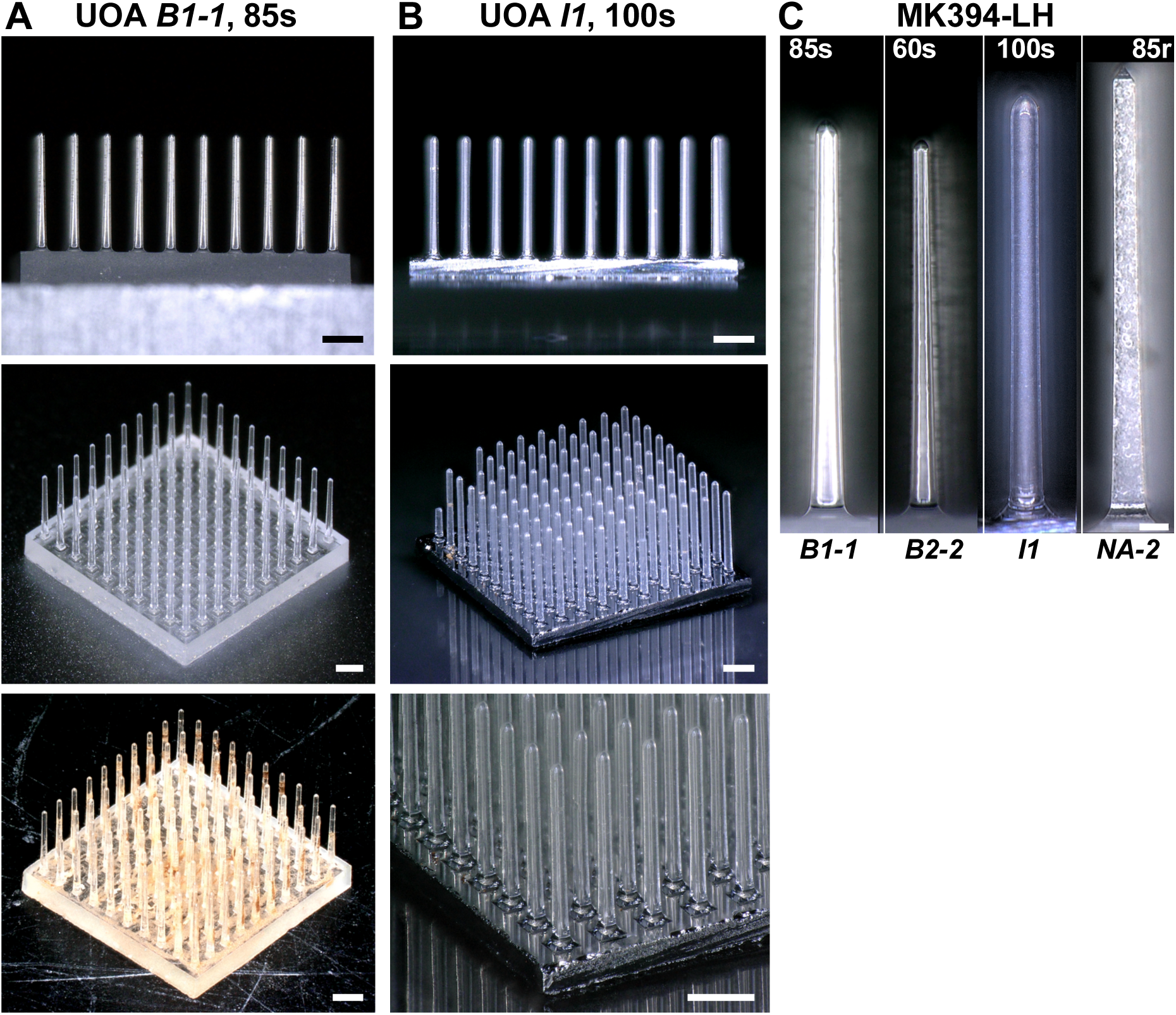
Implanted UOAs in case MK394-LH. **(A)** UOA *B1-1* (85µm shank diameter, 1.44mm shank length, smoot surface, round tip; see **Table 1**) before implantation viewed from the side (Top) and the top (Middle) and after explantation (Bottom). The explanted UOA retained all the shanks present prior to implantation. No shank was broken or damage as a result of implantation. **(B)** UOA *I1* (100µm shank diameter, 1.51mm shank length, smooth surface, round tip, with an optical interposer; see **Table 1**) before implantation viewed from the side (Top) and the top (Middle) and at higher magnification (Bottom). Prior to implantation this UOA had 4 damaged shanks, and a total of 96 intact shanks. Scale bars in (A-B): 500 µm. **(C)** Shank profile of each of the 4 UOAs implanted in case MK394-LH. Shank diameter and surface texture for each UOA are indicated at the top of each panel and case number at the bottom (for additional parameters see **Table 1**). Shank length for UOAs B2-2 and NA-2 was 1.37mm and 1.69mm, respectively. Scale bar: 100µm.

Recently, we developed and tested *in vivo* in macaque visual cortex^[28–29]^ a second generation device, which incorporates an optically opaque interposer layer with circular openings (or vias) in correspondence of the shanks, so that light emitted by the µLED array only transmits through the shank tips, preventing unwanted surface illumination via the inter-shank space and inter-shank cross talk. An example second generation device with an interposer layer is shown in **figure 1B** and in the bottom panel of **figure 2A** (the 100s UOA-*I1*). Here we assessed potential acute tissue damage caused by insertion of the UOA. To this goal we inserted a total of 16 “passive” UOAs (1 of which incorporated the interposer layer, *I1*) in one hemisphere of 3 macaque monkeys (**Fig. 2A**). The term “passive” refers to the UOA without integrated µLED arrays as opposed to the “active” UOA^[28–29]^. The implanted UOAs differed in shank diameter (60, 85, or 100 µm), surface texture (smooth, “s”, or rough, “r”) and tip geometry (round or sharp), as illustrated in **figure 1C**. Shank length varied across UOAs between 1.3 and 1.7 mm. For insertion, each UOA was positioned over the cortex and its backplane was struck with a high-speed pneumatic hammer specifically designed to minimize tissue damage for insertion of the Utah Electrode Array^[42]^ (see Experimental Methods). To minimize tissue damage from excessive pressure of the UOA backplane, we used a 1mm spacer, in order to obtain a partial insertion of the UOAs (all of which were longer than 1mm). To understand how insertion pressure affects tissue damage, we varied pulse pressure (9-20psi), while pulse duration was relatively constant across insertions. The UOAs were left in place for 1-2 hours (one was implanted for 3 hours) before being explanted and examined. An example UOA after explantation is shown in the bottom panel of **figure 1A**. The properties and insertion parameters of all implanted UOAs in each animal are reported in **Table 1**.

**Figure 2.**
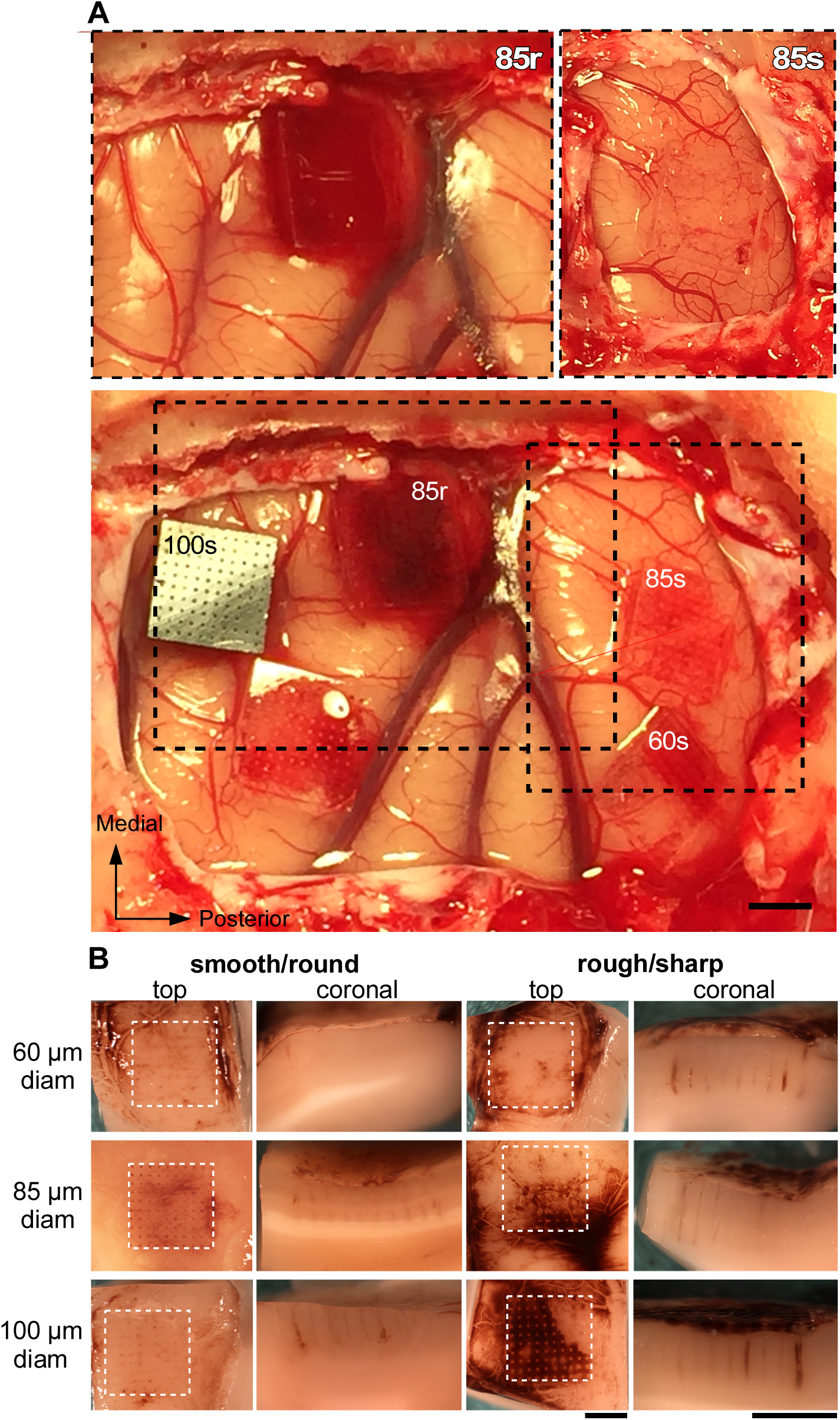
Surgical implantation of UOAs and macroscopic brain appearance after UOA explantation. **(A)** Five UOAs inserted in one hemisphere of a macaque monkey (case MK394-LH). Top: two UOAs of 85µm shank diameter, one with rough surface/sharp tip (85r; Left), the other with smooth surface/round tip (85s; Right) shown immediately after insertion. The 85r UOA caused significant bleeding immediately after insertion, while no bleeding was observed immediately after insertion of the 85s UOA. Bottom: all 5 UOAs on completion of the insertion (approximately 2 hours after insertion of the first UOA). *Dashed boxes:* same regions as shown in the top panels, 2 hours after insertion of the 85s UOA. The smooth/round UOAs caused less bleeding than the rough/sharp UOAs. There is more bleeding under the 85s UOA 2 hours after insertion compared to immediately after insertion (Top right). The 100s UOA has an interposer layer with optical vias. Scale bar: 2 mm, valid for all panels in (A). **(B)** Macroscopic examination of six example cortical implantation sites (one per UOA geometry), after UOA explantation. The left two columns show top and coronal views, respectively, of the implantation sites for the smooth/round UOAs, while the right two columns show the implantation sites for the rough/sharp UOAs. Top, Middle, and Bottom row: implantation sites of 60µm, 85µm, and 100µm shank diameter UOAs, respectively. Scale bars: 2mm valid for all panels in (B). All images in (B) are from case MK397-RH, except for the middle left two panels which are from case MK394-LH (**Table 1**).

**Table 1.** Properties and insertion parameters of UOAs.

| **CASE #** | **UOA#** | **Shank**<br>**Diameter**<br>**(μm)** | **Shank**<br>**Surface** | **Tip**<br>**geometry** | **Insertion**<br>**Pressure/Duration**<br>**(psi/ms)** | **Duration**<br>**of implant**<br>**(hrs:min)** | **Figures** |
| --- | --- | --- | --- | --- | --- | --- | --- |
| MK394-LH | B2-2 | 60 | smooth | round | 20/30 | 1:00 | 1C,2A |
|  | NA-2 | 85 | rough | sharp | 10/30 | 1:45 | 1C,2A |
|  | B1-1 | 85 | smooth | round | 10/30 | 2:00 | 1A,1C,2A |
|  | I1 | 100 | smooth | round | 15/30 | 1:20 | 1B-C,2A |
| MK397-RH | 60r | 60 | rough | sharp | 18/30 | 1:30 | 2B |
|  | 60s | 60 | smooth | round | 18/30 | 1:25 | 2B |
|  | 85r | 85 | rough | sharp | 18/30 | 1:50 | 2B |
|  | 85s | 85 | smooth | round | 18/30 | 1:45 | - |
|  | 100r | 100 | rough | sharp | 18/30 | 1:40 | 2B |
|  | 100s | 100 | smooth | round | 18/30 | 1:35 | 2B |
| MK406-RH | 2 | 60 | rough | sharp | 9/30 | 2:00 | 3B,5B,7B |
|  | 3 | 60 | smooth | round | 20/30 | 3:10 | 3A,5A,7A |
|  | 4 | 85 | rough | sharp | 9/50 | 1:30 | 3D,5D,7D |
|  | 5 | 85 | smooth | round | 9/50 | 1:50 | 3C,5C,7C |
|  | 6 | 100 | rough | sharp | 9/50 | 1:17 | 3F,5F,7F |
|  | 7 | 100 | smooth | round | 9/50 | 1:25 | 3E,5E,7E |

### 2.2. Macroscopic Examination

**Figure 2A** shows one example case (MK394-LH) in which 5 UOAs were inserted in the posterior half of one hemisphere in a macaque monkey. The top two panels show insertion of an 85µm-rough/sharp UOA (left panel) and an 85µm-smooth/round UOA (right panel) immediately after insertion, while the bottom panel shows all 5 UOAs on completion of the insertion (about 2 hours later). The rough surface UOA and those with largest shank diameters (100µm) typically caused more bleeding than the smooth and lower diameter UOAs, both immediately upon insertion and over the two-hour period.

**Figure 1B** shows the surface (top view) and a coronal view (across cortical layers) of the insertion sites after the UOA were explanted, for each type of inserted UOA (see **Table 1** for case numbers). Macroscopic examination of the cortical implantation site revealed that all UOAs caused minimal brain edema. In some cases, we observed interstitial microhemorrhages emanating from the tracks left by the optrode shanks that extended in one or more directions. These microhemorrhages were limited to within a few millimeters of the optrode tracks and were more evident at the sites of rough/sharp UOA insertion compared to smooth/round UOA insertion sites. This damage seemed to result from the blood vessels encountered in the path of the penetrating optrode shanks, in addition to some mechanical damage of the small capillaries.

### 2.3. Microscopic Examination: Glial Response and Neuronal Viability

#### 2.3.1. Astrocyte Activation

To identify both resting and activated astrocytes, we immunostained coronal tissue sections for the glial fibrillary acidic protein (GFAP). For each implanted UOA, we selected one tissue section, specifically the one containing the largest number of tracks left by the inserted device shanks and full shank lengths. On this section, we quantified the amount of GFAP immunostaining as Integrated Density (see Experimental Methods for how this was measured), at 4 depths along each of the tracks left by the UOA shanks, specifically at 0-200 µm, 400-600 µm, 800-1000 µm and 1300-1500 µm, approximately corresponding to cortical layers 1-2, 3, 4 and 5-6, respectively. Figure 3A-B shows representative images of GFAP staining at the UOA implantation site and in control non-implanted tissue from the same tissue section, for a smooth (A) and a rough (B) UOA of 60µm shank diameter. Figure 3C-D **and 3E-F** show equivalent images for UOAs of 85µm and 100µm shank diameter, respectively.

**Figure 3.**
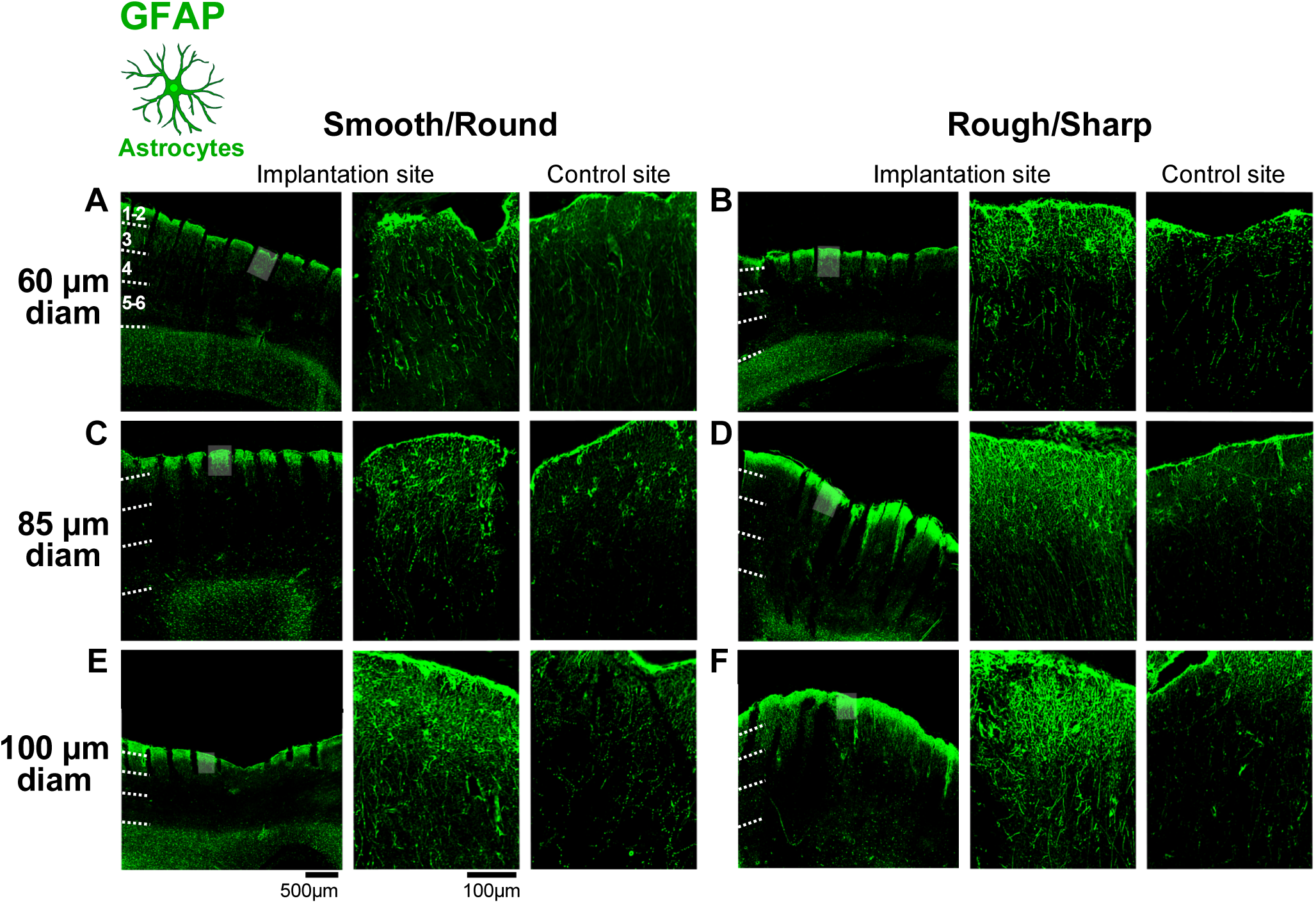
GFAP immunoreactivity in response to insertion of UOAs of different shank diameters and geometries. **(A)** Left: low power micrograph of a coronal section immunostained for GFAP at the site of implantation of a 60µm-shank diameter smooth/round UOA. The tracks left by the UOA shanks are visible. The area corresponding to the *shaded box* is shown at higher power in the middle panel. *Dashed* lines mark cortical layer boundaries, which are indicated. Middle: higher magnification of the region corresponding to the shaded box in the left panel. Right: higher magnification micrograph of GFAP-IHC at a non-implanted control site in the same section. **(B)** Same as in (A), but for a rough/sharp UOA implantation and respective control sites. **(C)** Same as (A), but for an 85µm-shank diameter smooth/round UOA implantation and respective control sites. **(D)** Same as (B), but for an 85µm-shank diameter rough/sharp UOA implantation and respective control sites. **(E)** Same as (A,C), but for a 100µm-shank diameter smooth/round UOA implantation and respective control sites. Scale bars: 500µm (left) and 100µm (right), and valid for all equivalent panels in (A-F). **(F)** Same as (B,D), but for a 100µm-shank diameter rough/sharp UOA implantation and respective control sites. Case number is: MK406RH for panels A-F (**Table 1**).

GFAP immunoreactivity was increased, relative to control sites, near the shank tracks, primarily in the most superficial layers (L1-2), forming a densely packed layer of astrocytes across all evaluated UOA insertions. Control sites, instead, showed a normal population of GFAP+ astrocytes. Qualitatively, GFAP immunoreactivity appeared to increase with UOA shank diameter, and for each diameter it was higher for rough vs. smooth UOAs (Fig. 3).

In figure 4A, GFAP+ immunostaining is quantified at each cortical depth for UOAs of different diameters and geometries (smooth/round vs. rough/sharp), as GFAP Integrated Density at the implantation site normalized to GFAP Integrated Density at each respective control site. Note that in the figures, for simplicity we use the layer nomenclature, but measurements were made at fixed depths across cortical areas of UOA insertions. Overall, GFAP immunoreactivity increased with increasing UOA shank diameter and roughness of the shanks. Thus, at all cortical depths both larger diameters and rough/sharp geometries generally, although not always, elicited more significant astrocyte reactions. **Supporting** Figure 1 shows the results of the statistical comparisons. Specifically, in **Supporting** Figure 1A for each UOA shank diameter, raw GFAP Integrated Density for smooth and rough UOAs is compared to GFAP Integrated Density in the control, across cortical depths (Mann-Whitney test). In all layers, most UOAs, irrespective of shank diameter and smooth or rough surface, caused a significant increase in GFAP immunoreactivity compared to control, but this increase was generally greater, and more significantly so, for the rough/sharp UOA geometry. **Supporting** Figure 1B shows how shank diameter affects astrocyte response. In this figure, absolute GFAP Integrate Density for smooth/round UOAs and rough/sharp UOAs is compared across UOAs of different shank diameters (Kruskal-Wallis test, with Dunn’s correction for multiple comparisons). For the smooth/round UOAs, larger diameters caused significantly greater increases in GFAP immunoreactivity in the layers of UOA insertion, i.e. L1-4, but no significant differences in GFAP immunoreactivity were observed in deeper tissue (L5-6) across UOA of different diameters. For the rough/sharp UOAs, in all layers the 85µm shank diameter ones caused the largest increase in GFAP immunoreactivity, while the 100µm shank diameter UOAs caused the least increase in GFAP immunoreactivity at all depths, except in L1-2. **Supporting** Figure 1C shows how surface texture affects astrocyte response. For each UOA shank diameter, GFAP Integrated Density is compared for smooth/round vs. rough/sharp UOAs across cortical depths (Mann-Whitney test). At the smaller diameters (60 and 85µm), the rough/sharp UOAs generally caused greater increases in GFAP immunoreactivity compared to smooth/round UOAs. For the 100µm diameter UOAs, either there was no difference in GFAP Integrated Density between smooth/round and rough/sharp devices or the latter showed lower GFAP Integrated Density. This suggests that the largest diameter UOAs cause significant damage whether they are smooth/round or rough/sharp, but it is possible that at this larger shank diameter, the sharper tip may cause less damage, as it penetrates tissue more easily than the round tip (see Discussion).

**Figure 4.**
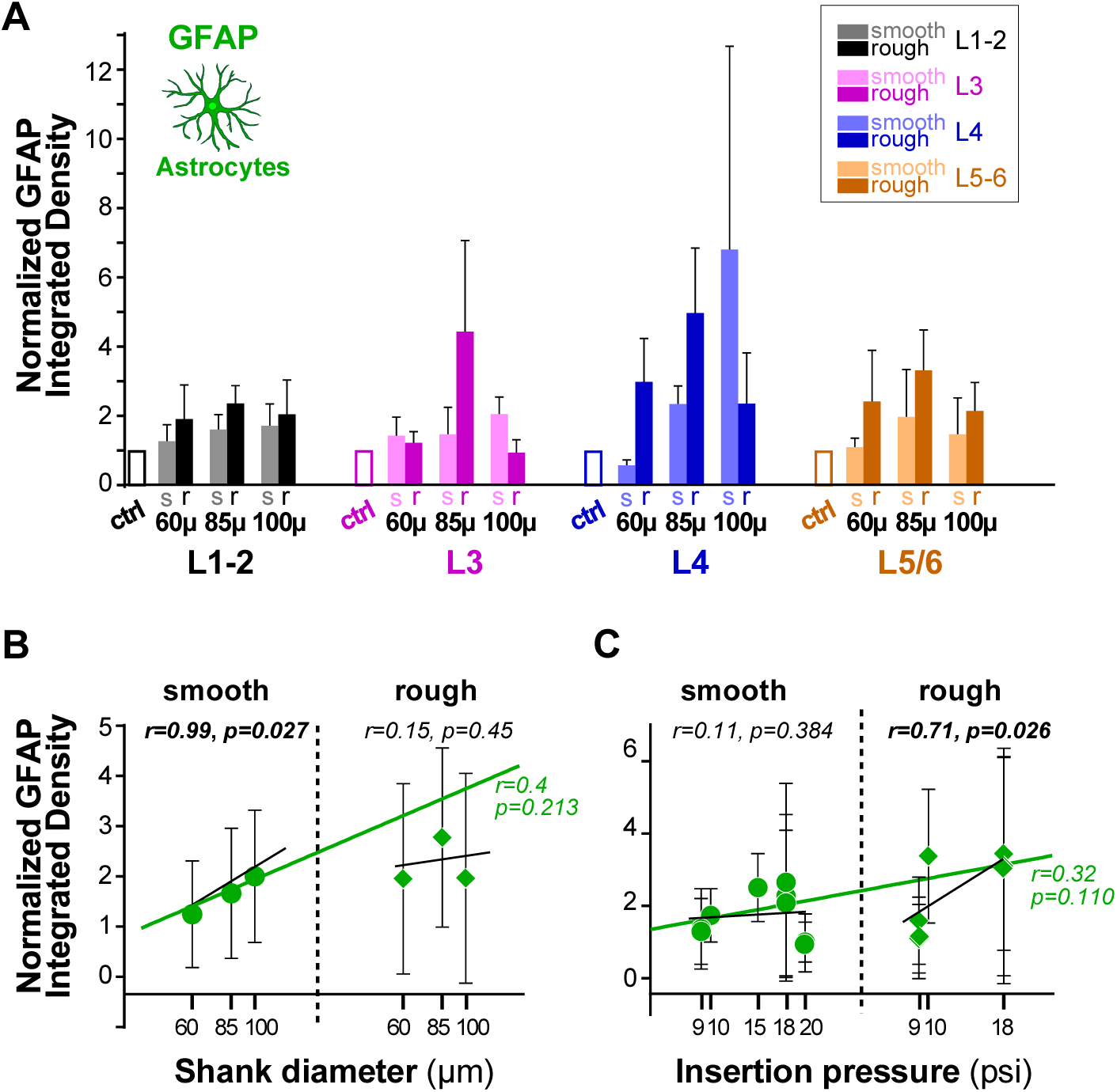
Quantification of GFAP immunoreactivity across cortical depths/layers in response to UOA insertion. (**A**) Integrated GFAP Density (a measure of astrocyte activation, see Experimental Methods) at different cortical depths (here indicated as cortical layers) following implantation of UOAs of different diameters (60,85,100 µm) and geometries (*s:* smooth/round; *r:* rough/sharp). Integrated Density at the implantation site is normalized to Integrated Density at the control site (*ctrl*). Error bars: s.e.m. (**B**) Normalized GFAP Integrated Density as a function of shank diameter for smooth/round (*circles*) and rough/sharp (*diamonds*) UOAs. *Black lines:* lines of best fit (regression) for the individual smooth and rough UOA populations. *Green line:* line of best fit for the two populations pooled together. r and p values are indicated. (**C**) Normalized GFAP Integrated Density as a function of pressure applied with the pneumatic inserter to insert the UOA into the cortex. Other conventions are as in (B).

**Figure 4B** summarizes how GFAP immunoreactivity varies as a function of UOA shank diameter separately for smooth/round and rough/sharp UOAs, as well as jointly (smooth and rough pooled together by shank diameter). Overall, we observed a positive correlation between GFAP Integrated Density and shank diameter for both geometries, although this correlation was statistically significant only for the smooth/round geometry (r=0.99, p=0.0268, Pearson’s correlation).

We also investigated how GFAP immunoreactivity varied as a function of the pressure applied with the pneumatic inserter during UOA insertion. We found a statistically significant positive correlation between GFAP Integrated Density and insertion pressure only for the rough/sharp geometry (r=0.71, p=0.0257, Pearson’s correlation; Fig. 4C).

#### 2.3.2. Microglial Activation

To identify resting and activated microglia/macrophages, we quantified immunostained tissue sections for the calcium binding adaptor molecule, Iba1, at the same 4 depths along the UOA shank tracks used for GFAP analysis. Representative images of Iba1 immunohistochemistry (IHC) at the implantation and control sites for UOAs of various diameters and geometries are shown in figure 5. This analysis was performed on the same sections used for the analysis of GFAP immunoreactivity as these sections were double-stained for both GFAP and Iba1 (see **Supporting** Fig. 2). For all cases qualitative observations indicated increased Iba1 immunoreactivity compared to control sites, and this increase appeared greater at the 100µ-diameter UOA implantation sites, particularly for the rough/sharp geometries.

**Figure 5.**
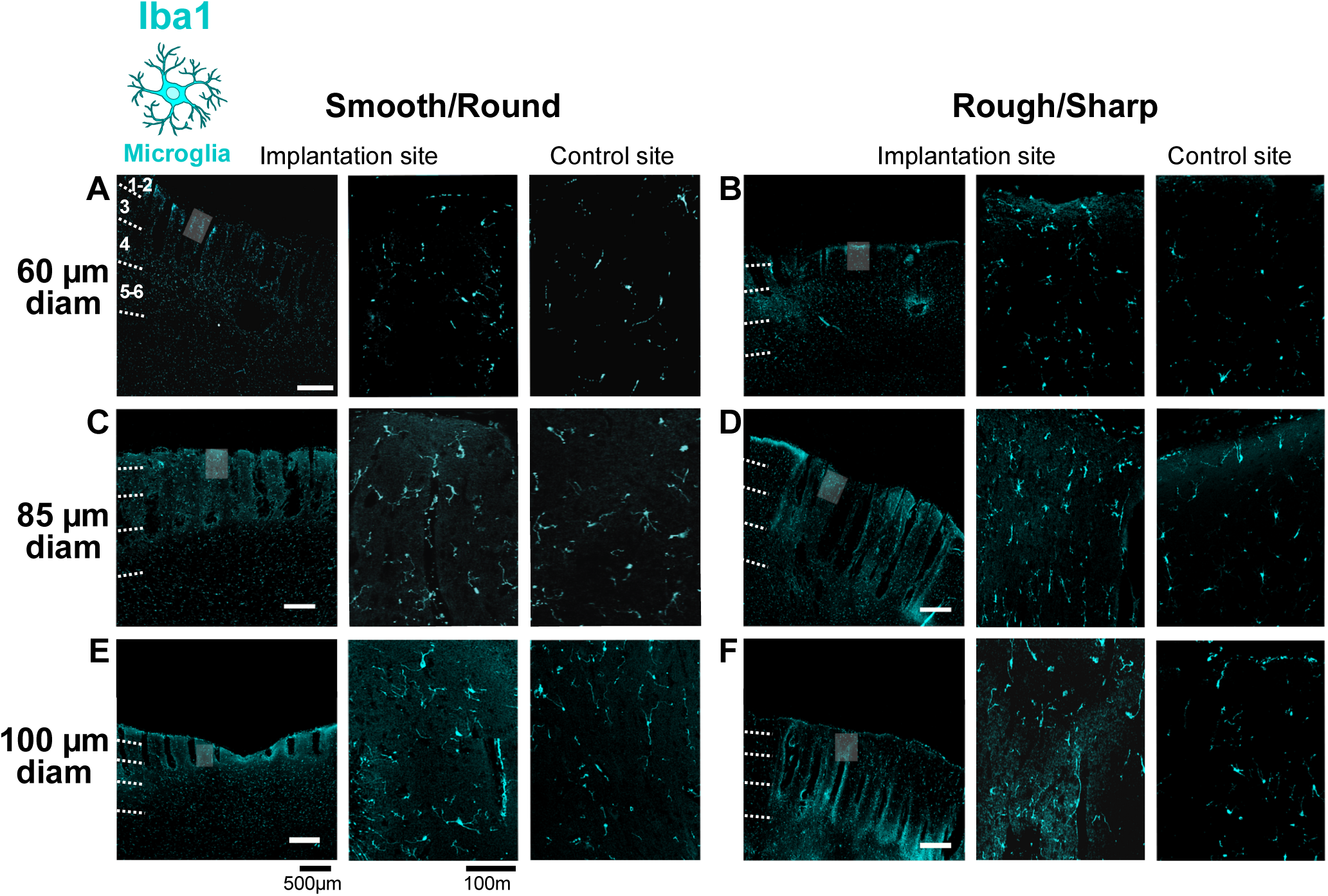
Iba1 immunoreactivity in response to insertion of UOAs of different shank diameters and geometries. **(A)** Left: low power micrograph of a coronal section immunostained for Iba1 at the site of implantation of a 60µm-shank diameter smooth/round UOA. Conventions are as in **Fig. 3A**. Middle: higher magnification of the region corresponding to the shaded box in the left panel. Right: higher magnification micrograph of Iba1-IHC at a non-implanted control site in the same section. **(B)** Same as in (A), but for a rough/sharp UOA implantation and respective control sites. **(C)** Same as (A), but for an 85µm-shank diameter smooth/round UOA implantation and respective control sites. **(D)** Same as (B), but for an 85µm-shank diameter rough/sharp UOA implantation and respective control sites. **(E)** Same as (A,C), but for a 100µm-shank diameter smooth/round UOA implantation and respective control sites. Scale bars: 500µm (left) and 100µm (right), and valid for all equivalent panels in (A-F). **(F)** Same as (B,D), but for a 100µm-shank diameter rough/sharp UOA implantation and respective control sites. Case number is: MK406RH for panels A-F (**Table 1**).

In figure 6A, Iba1+ immunostaining is quantified across cortical depths/layers for UOAs of different diameters and geometries (smooth/round vs. rough/sharp), as Iba1 Integrated Density at the implantation site normalized to Iba1 Integrated Density at each respective control site, and statistical comparisons are shown in **Supporting** Figure 3. Iba1 immunoreactivity was increased relative to controls at all shank diameters at the most superficial depths (L1-2), but in deeper layers only the 100µm-diameter UOAs caused significant increases relative to controls (**Supporting** Fig. 3A). The 100µm diameter UOAs caused a higher increase in Iba1 immunoreactivity compared to the 60µm and often the 85µm diameter UOAs, but in most instances there was no significant difference in Iba1 immunoreactivity at the implantation sites of the 60µm and 85µm diameter UOAs (**Supporting** Fig. 3B). Shank geometry did not seem to significantly affect Iba1 immunoreactivity, but in some layers, particularly for the smallest diameter UOAs, the smooth/round geometry caused slightly higher Iba1 immunoreactivity compared to the rough/sharp geometry. This difference was statistically significant for the 60µm shank diameter in L1-2 (*p=0.0267*) and L4 (p=0.0358), and for the 85µm shank diameter in L3 (*p<0.0001*) and L5/6 (*p<0.0144*) (**Supporting** Fig. 3C). Figure 6B summarizes the overall effect of diameter on Iba1 immunoreactivity. There was a strong positive correlation between Iba1 Integrated Density and shank diameter for both smooth (r=0.99; Pearson correlation) and rough (r=0.96) geometries, as well as for both geometries pooled together (r=0.9), but this correlation was only statistically significant for the smooth geometry (p=0.0308) and the overall population pooled across geometries (p=0.0071). Insertion pressure did not affect Iba1 immunoreactivity; there was no significant correlation between Iba1 immunoreactivity and UOA insertion pressure (Fig. 6C).

**Figure 6.**
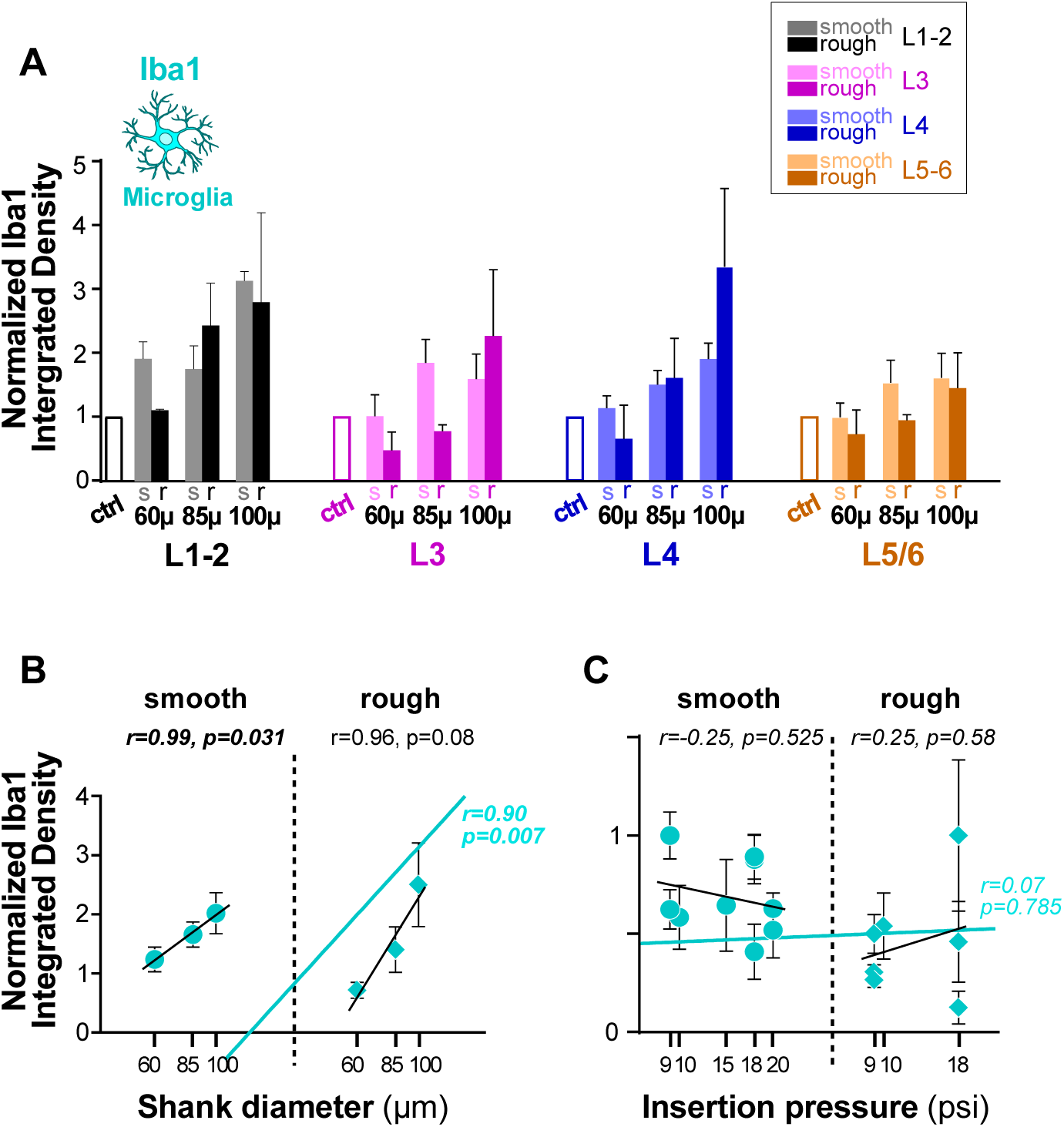
Quantification of Iba1 immunoreactivity across cortical depths/layers in response to UOA insertion. **(A)** Integrated Iba1 Density (a measure of microglia activation) in different layers following implantation of UOAs of different diameters and geometries (*s:* smooth/round; *r:* rough/sharp). Integrated Density at the implantation site is normalized to Integrated Density at the control site (*ctrl*). Error bars: s.e.m. **(B)** Normalized Iba1 Integrated Density as a function of shank diameter for smooth/round (*circles*) and rough/sharp (*diamonds*) UOAs. *Black lines:* lines of best fit (regression) for the individual smooth and rough UOA populations. *Cyan line:* line of best fit for the two populations pooled together. r and p values are indicated. **(C)** Normalized Iba1 Integrated Density as a function of pressure applied with the pneumatic inserter to insert the UOA into the cortex. Other conventions are as in (B).

#### 2.3.3. Neuronal viability

Survival of neurons is crucial for the success of technologies based on optogenetics. Furthermore, neuronal cell bodies must be as close as possible to the optrode shanks for optimal device function. We used IHC against the neuronal nuclear protein NeuN to identify neurons. This protein is localized in the nuclei and perinuclear cytoplasm of most types of neurons across the nervous system and is not expressed in glial cells, therefore it is a specific neuronal marker. Representative images of NeuN-IHC at the implantation and control sites for UOAs of various diameters and geometries are shown in figure 7. This analysis was performed on the same sections used for the analysis of GFAP immunoreactivity, which were double-stained for both GFAP and NeuN (see **Supporting** Fig. 4). Qualitative observations indicated that for UOAs of smooth/round geometry, NeuN Integrated Density was reduced for the 85µm and 100µm-diameter UOAs relative to control, but not for the smallest diameter UOA. Moreover, for this geometry, NeuN Integrated Density appeared to decrease with increasing UOA diameter (Fig. 7A**,C****,E**). For the rough/sharp UOA geometry, instead, all, but the 100µm diameter, UOAs caused a decrease in NeuN immunoreactivity relative to control. Surprisingly NeuN Integrated Density appeared higher for the 100µm diameter rough/sharp UOA compared to both the control and implantation sites of smaller diameter UOAs (Fig. 7B**,D****,F**).

**Figure 7.**
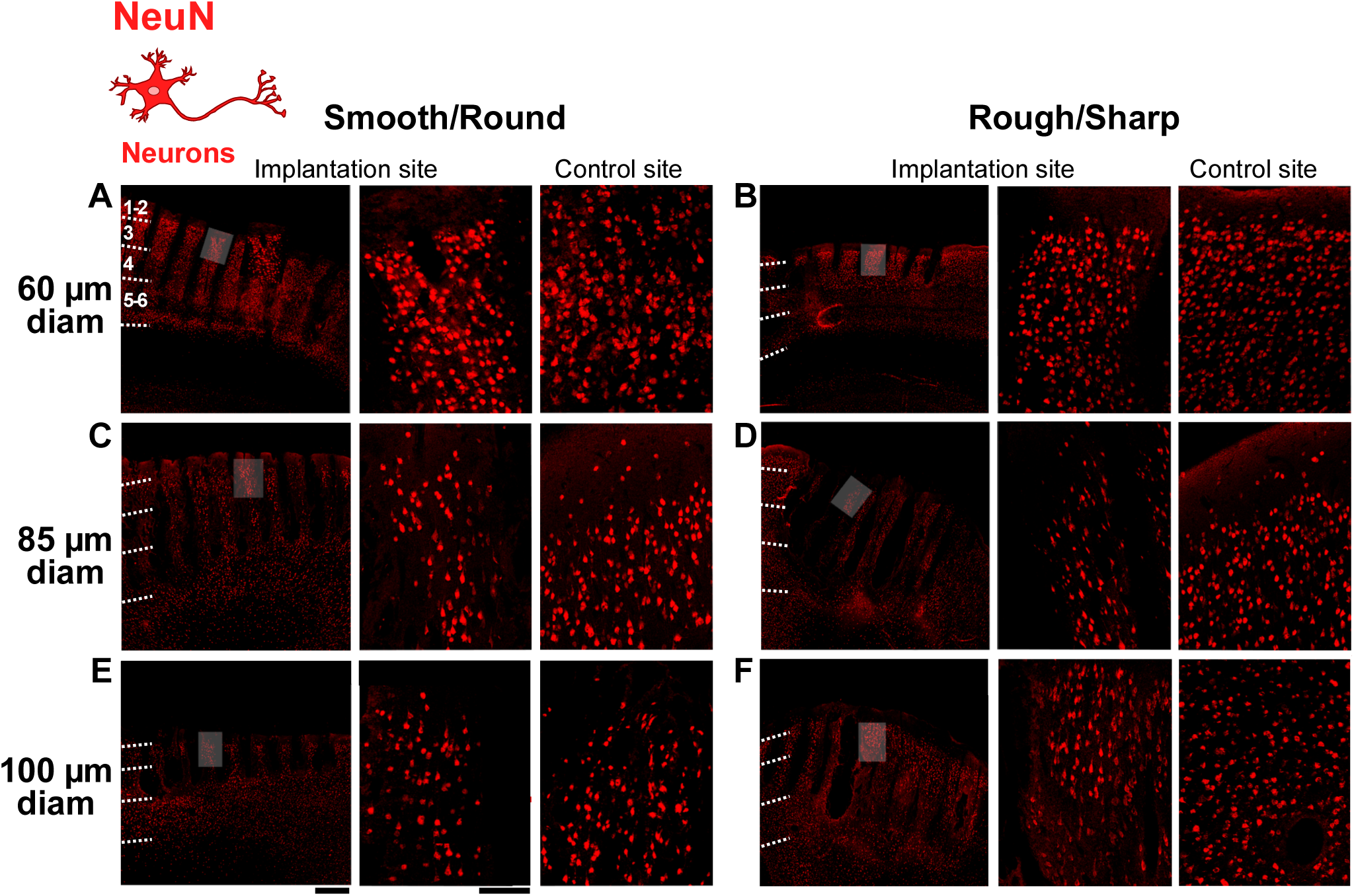
NeuN immunoreactivity in response to insertion of UOAs of different shank diameters and geometries. **(A)** Left: low power micrograph of a coronal section immunostained for NeuN at the site of implantation of a 60µm-shank diameter smooth/round UOA. Conventions are as in **Fig. 3A**. Middle: higher magnification of the region corresponding to the shaded box in the left panel. Right: higher magnification micrograph of NeuN-IHC at a non-implanted control site in the same section. **(B)** Same as in (A), but for a rough/sharp UOA implantation and respective control sites. **(C)** Same as (A), but for an 85µm-shank diameter smooth/round UOA implantation and respective control sites. **(D)** Same as (B), but for an 85µm-shank diameter rough/sharp UOA implantation and respective control sites. **(E)** Same as (A,C), but for a 100µm-shank diameter smooth/round UOA implantation and respective control sites. Scale bars: 500µm (left) and 100µm (right), and valid for all equivalent panels in (A-F). **(F)** Same as (B,D), but for a 100µm-shank diameter rough/sharp UOA implantation and respective control sites. Case number is: MK406RH for panels A-F (**Table 1**).

In figure 8A, NeuN immunostaining is quantified across cortical depths for UOAs of different diameters and geometries (smooth/round vs. rough/sharp), as NeuN Integrated Density at the implantation site normalized to NeuN Integrated Density at each respective control site, and statistical comparisons are shown in **Supporting** Figure 5. Implantation of smooth/round UOAs of 85µm and 100µm diameters significantly decreased NeuN immunoreactivity relative to controls in most layers. The 60µm diameter smooth/round UOAs, instead, caused no change in NeuN Integrated Density relative to controls in all layers, but L1/2 where they showed a slight increase in NeuN Integrated Density relative to the control. The rough/sharp UOAs of 60µm and 85µm diameter significantly reduced NeuN Integrated Density in most layers compared to controls. In contrast, and consistent with qualitative observations, the 100µm diameter UOAs of rough/sharp geometry caused no significant change in NeuN Integrated Density relative to controls in all layers, except L1-2, where NeuN Integrated Density was increased by the largest diameter rough UOAs (**Supporting** Fig. 5A). These slight increases in NeuN immunoreactivity relative to control sites are likely due to tissue compression around the optrode shanks caused by UOA insertion. The effect of diameter on NeuN immunoreactivity is shown in **Supporting** figure 5B. For the smooth/round geometry, NeuN Integrated Density was significantly lower for the larger diameter UOAs compared to the 60µm diameter UOA. In contrast, for the rough/sharp geometry, NeuN Integrated Density was higher for the 100µm diameter UOAs compared to the smaller diameter UOAs. Shank geometry had opposite effects for small vs. large diameter UOAs, with rough/sharp geometries being associated with lower NeuN Integrated Density in all layers for the 60µm diameter UOA and in L1-2 for the 85µm diameter UOA, but it was associated with higher NeuN Integrated Density for the 100µm diameter UOA (**Supporting** Fig. 5C). A similar result is conveyed in Figure 8B, which shows a high negative correlation (r=0.87) of NeuN Integrated Density with shank diameter for smooth UOAs, but a positive correlation (r=0.96) for rough/sharp UOAs (albeit these correlations did not reach statistical significance). NeuN Integrated Density across the population of smooth and rough UOAs showed a statistically significant positive correlation (r=0.63, p=0.0103; Pearson correlation) with UOA insertion pressure (Fig. 8C), suggesting increasing tissue compression with insertion pressure and consequent apparent increase in neuronal density. This phenomenon may be the cause for the apparent increase in NeuN Integrated Density that we observed for the largest diameter rough UOAs. It is also possible that for the largest diameter UOAs the round tip causes relatively more damage (greater neuronal death) than the sharp tip as noted for other IHC markers above.

**Figure 8.**
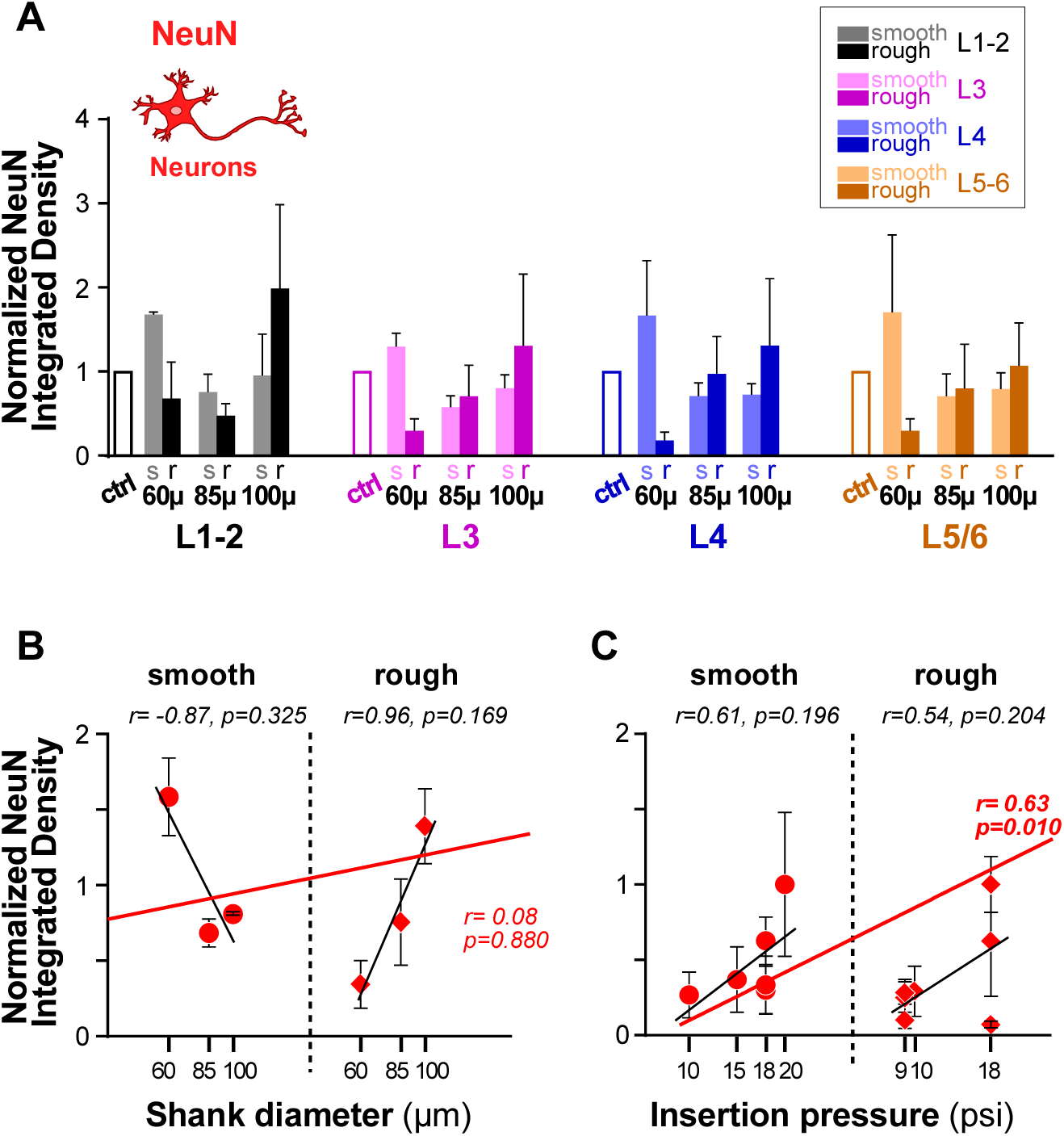
Quantification of NeuN immunoreactivity across cortical depths/layers in response to UOA insertion. (**A**) Integrated NeuN Density (a measure of neuron density) in different layers following implantation of UOAs of different diameters and geometries (*s:* smooth/round; *r:* rough/sharp). Integrated Density at the implantation site is normalized to Integrated Density at the control site (*ctrl*). Error bars: s.e.m. (**B**) Normalized NeuN Integrated Density as a function of shank diameter for smooth/round (*circles*) and rough/sharp (*diamonds*) UOAs. *Black lines:* lines of best fit (regression) for the individual smooth and rough UOA populations. *Red line:* line of best fit for the two populations pooled together. r and p values are indicated. (**C**) Normalized NeuN Integrated Density as a function of pressure applied with the pneumatic inserter to insert the UOA into the cortex. Other conventions are as in (B).

## 3. Discussion

In this study, we have identified key factors in designing Optrode Arrays for neural applications. We found that the factors that most affect tissue damage are shank diameter, surface texture and tip geometry. Larger shank diameters cause greater tissue damage, leading to higher astrocytic and microglial activation, and reduced neuronal viability. Smooth-texture, round-tipped shanks help reduce astrocytic activation and preserve neuronal viability for the smaller shank diameter UOAs (60, 85 µm), but for the largest (100µm) shank diameter UOAs, the round tip can be more detrimental than a rough surface texture. Higher insertion pressures have limited effects on the inflammatory response, but lead to greater tissue compression. Therefore, balancing shank diameter, tip geometry, and insertion pressure is essential for an effective, tissue-friendly UOA design.

### 3.1. Influence of Shank Diameter

UOAs with shanks of 60µm diameter, smooth texture and round tips caused minimal damage. Compared to normal control tissue, tissue implanted with these UOAs showed a mild increase in astrocytic and microglia activation only in the most superficial cortical layers, and no significant reduction in neuronal viability. However, larger shank diameters significantly influenced astrocytic and microglial activation, as indicated by elevated GFAP and Iba1 immunoreactivity around the implantation sites. This effect is likely due to the greater physical disruption of tissue caused by larger diameter shanks, potentially affecting a wider area of cells and capillaries, which is consistent with previous research findings^[43–45]^. This mechanical disruption induces a proinflammation cascade, causing the mobilization of astrocytes and microglia to the insult site to manage the tissue injury and begin the repair process^[33]^. At the same time, however, larger diameter shanks allow for more efficient light coupling between the µLEDs and the shanks for optogenetic applications^[46]^, and are easier to manufacture. Furthermore, increased light coupling efficiency leads to less heat generation during μLED operation. We previously demonstrated effective light delivery to deep cortical layers of macaque visual cortex *in vivo* using 100µm-shank diameter active UOAs^[28–29]^. However, the results of the present study suggest that UOAs of smaller shank diameter could be of benefit in reducing the acute inflammatory response as well as confining it to the region of insertion. Ultimately, this represents a trade-off in the balance between achieving optimal light transmission while preserving tissue integrity.

Neuronal viability also varies with shank diameter and surface texture. For the smooth/round UOAs, the larger diameter shanks reduced NeuN immunoreactivity compared to the smaller diameter shanks, indicating reduced neuronal preservation. Notably, for the rough/sharp UOAs, instead, the largest diameter shanks were associated with higher NeuN immunoreactivity compared to the smaller and intermediate diameter shanks. While on the surface this may seem to indicate better neuronal preservation, we suspect that this effect likely depends on factors beyond diameter alone. In particular, because NeuN immunoreactivity following implantation of large diameter, rough/sharp UOAs was increased relative to controls, it is likely that the apparent increase in NeuN immunoreactivity is the result of greater tissue compression caused by insertion of larger diameter rough UOAs.

### 3.2. Influence of Surface Texture and Tip Geometry

A well-designed probe can greatly improve long-term biocompatibility and biotolerability. Optimizing both shank and tip biocompatibility reduces tissue damage and enhances their overall performance^[47]^. While common features, such as the relationship between device design— namely, shape, size, and tethering—and its long-term stability, have been considered in previous research^[48]^, the effect of shank surface texture on the tissue-shank interface remains poorly understood. Our results show that smooth shanks in general caused the least disruption to the surrounding cortical tissue, but this depended on shank diameter. The smooth-surface/round-tip geometry for smaller diameter shanks minimized astrocytic activation and neuronal loss, while microglia appeared to be less significantly influenced by tip geometry and shank surface texture, although there was a tendency for microglia activation to be minimized by the rough/sharp geometry of smaller shank diameters. The smooth surface likely reduces friction during insertion, thereby minimizing mechanical trauma and inflammatory responses. Additionally, the round tip of small diameter shanks enables a gentler insertion path, localizing stress to surrounding cells. While the sharp tip facilitates penetration, it may also elevate mechanical stress on surrounding brain cells. Overall, these findings suggest that for smaller shank diameters, smooth shank surfaces and round tips effectively reduce astrocytic activation and certain inflammatory responses, and better preserve neuronal viability; however, they do not completely mitigate microglial activation, which appears to favor rough surfaces/sharp-tipped shanks. In contrast to these results, the rough/sharp geometry minimized astrocytic and microglia activation in response to implantation of UOAs of larger, 100µm, diameter shanks. It is likely that for the wider shanks the sharp tip facilitates penetration, thus causing less damage than the thicker round tip. These results underscore the need for a delicate balance in UOA design for optimal biocompatibility.

Neuronal viability results indicate that smaller shank diameters with smooth surfaces are more effective in preserving neuronal density compared to larger, rougher shanks. Interestingly, larger shanks with rough/sharp geometry show relatively increased neuronal density, likely due to increased tissue compression, as well as reduced insertion resistance (due to the sharper tip), which minimizes physical disruption and neuronal death around the site. This suggests that while smaller diameters generally tend to be less injurious, incorporating a sharp tip into a larger diameter may partially counteract the adverse effects typically associated with increased shank size, if increased shank size is used for light coupling reasons. These results are consistent with those of a recent study^[41]^, which showed that for microwire diameters of <80µm, tip shape has little effect on insertion force, but for microwire diameters of about 120µm, sharp tips have lower insertion forces than rounded tips, hence causing less damage. The same study also showed that sharper tips produce less vascular damage, as do smaller diameter microwires.

### 3.3. Influence of Insertion Pressure

UEAs traditionally require a minimum insertion speed of 8.3 m/s to ensure the full, safe insertion of all 100 electrodes in the array to a depth of 1.5 mm at a pressure of 25 to 29 psi^^[42, 49]^^. For UOAs, a high insertion speed is also required, with previous reports using insertions pressures of 20 psi and a pulse width of 30 ms^[28–29]^. The present study tested the implantation of UOAs at high insertion speeds and pressures ranging from 9 to 20 psi with a pulse duration of either 30 or 50 ms. Results indicate that higher insertion pressures lead to greater mechanical stress on the tissue, as shown by higher astrocytic activation for rough designs, while microglia seemed to be unaffected by insertion pressure. Effects on neuronal viability suggest that increased insertion pressure leads to greater tissue compression, and, therefore, to an apparent increase in neuronal density.

## 4. Conclusion

In summary, this research highlights the critical importance of achieving a balanced design for UOAs that prioritizes effectiveness, reliability, and safety. Optimizing surface and tip designs alongside controlled insertion pressures enhances UOA tissue-stability while preserving tissue integrity, establishing a foundation for improved long-term performance in both optogenetic and neurostimulation applications. UOAs with shanks of smaller diameter, smooth surface texture and round tips cause the least damage, but maybe less efficient in delivering light to deeper tissue than larger diameter UOAs. On the other hand, 100µm diameter UOAs have been previously shown to effectively photoactivate deep cortical tissue *in vivo* without compromising neuronal responsiveness, causing tissue damage comparable to, or even lesser, than that caused by the widely-used, FDA-approved UEAs. Future studies should focus on understanding the long-term effects of UOA implantation, as these may differ from acute responses. Complementary research is also needed to explore coatings or materials that reduce friction and inflammatory responses, enabling safer implantations.

## 5. Experimental Methods

### Experimental Design

For each experiment, four to six “passive” UOAs (lacking integrated µLED arrays) were acutely implanted in the cerebral cortex of one hemisphere of anesthetized macaque monkeys. On completion of the insertion, the animals were euthanized and perfused with fixative. The brains were processed for histology and IHC to identify markers of inflammation and neuronal death, and immunohistochemical markers were analyzed quantitatively.

### Animals

Three adult female Cynomolgus monkeys (*Macaca fascicularis*) were used in this study (see **Table 1**). All procedures adhered to the guidelines outlined in the National Institutes of Health Guide for the Care and Use of Laboratory Animals and received approval from the University of Utah Institutional Animal Care and Use Committee (IACUC).

### Surgical Procedures

Implantation of passive UOAs was performed on the last day of an unrelated terminal electrophysiological recording experiment performed over a period of 4-5 days on the hemisphere contralateral to the implanted one. At the time of UOA implantation the animals were placed in a stereotaxic apparatus, and had been maintained under anesthesia for several days by continuous infusion of sufentanil citrate (5-10 µg/kg/h). Animals were artificially ventilated with 100% oxygen, and vital signs (heart rate, end tidal CO2, oxygen saturation, electrocardiogram, and body temperature) were continuously monitored for the duration of the experiment. I.V. fluids were delivered at a rate of 3 cc/kg/h. Following scalp incision, a large craniotomy and durotomy were made encompassing all of the visual cortex, and parts of the auditory, motor, and somatosensory cortices, to allow space for 4-6 device implantations (see e.g. Fig. 2A). The UOAs were positioned over the cortex, and then inserted, by a neurosurgeon (J.D.R.) using a high-speed pneumatic hammer designed to minimize tissue damage during insertion of the Utah Electrode Array ^[42]^ (Electrode Inserter System - Blackrock Neurotech, Salt Lake City, UT). On completion of the insertions, the animals were sacrificed with Beuthanasia (0.22 ml/kg, i.p.) and perfused transcardially with saline for 2-3 minutes, followed by 4% paraformaldehyde (PFA) in 0.1 M phosphate buffer for 20-25 minutes.

### UOA Insertion

A total of 16 10×10 passive UOAs were implanted in 3 animals, 5-6 in one hemisphere of each animal. The inserted UOAs differed in several parameters (**Table 1**), including, shank diameter, surface texture and tip geometry. Shank length varied across UOAs between 1.3 to 1.7 mm. We also varied insertion parameters, primarily pulse pressure, while pulse duration was relatively constant across insertions (the dial was set at 30 or 50 a.u.). Pressure and pulse settings were first calibrated by testing against a gloved finger that device engagement resulted in a single strike of the insertion hammer. Moreover, to ensure a clean delivery of the UOA into the cortex with no pullback of the UOA during retraction of the hammer (due to surface tension at the hammer/UOA interface), a drop of sterile saline was placed on a thin periosteal elevator, the elevator was gently placed against the backplane of the UOA and then struck with the insertion hammer. To minimize tissue damage from excessive pressure of the UOA backplane, in these experiments we used a 1mm spacer, in order to obtain a partial insertion of the UOAs, all of which had shank lengths >1mm.

### Histology and Immunohistochemistry

After perfusion, each brain was carefully extracted from the skull. The UOAs were explanted from the cortex and examined. The brain was post-fixed in fresh fixative (4% PFA) for 2 days. Each UOA-implanted cortical site was blocked from the rest of the brain, and each individual block was hemisected to facilitate clear visualization of the optrode shanks (as shown in Fig. 2B). Prior to sectioning, each block was cryoprotected by equilibrating in a step-wise gradient of sucrose (15%, 20%, and 30%). Each block was then frozen at −25 °C in Tissue-Tek, and coronally sectioned at a thickness of 20µm using a cryostat (HM505E, Microm). Tissue sections were mounted onto glass slides and stored at −80°C until immunohistochemical labeling.

For IHC, tissue sections were first equilibrated to room temperature (RT) to allow for optimal adhesion to microscope slides. The sections were then washed three times in phosphate buffered saline (PBS; pH 7.4) for 10 minutes and incubated in bovine serum albumin (BSA, 10%, Sigma, St. Louis, MO; Millipore CAS#9048-46-8) in PBS containing 0.5% Triton-X (PBS-T) (Sigma) for one hour at RT before being incubated for 24 hours at 4°C with primary antibodies diluted in 2% BSA + 0.5% PBS. The primary antibodies employed for staining astrocytes, neurons, and microglia were: chicken polyclonal anti-GFAP (1:200; Ab5541 Millipore, Germany; RRID:AB_177521), rabbit polyclonal anti-NeuN (1:200; AbN78. Millipore, Germany; RRID:AB_10807945), and rabbit polyclonal anti-Iba1 (1:100; Ab178846, Abcam, UK; RRID:AB_2636859), respectively. The sections were then washed, and incubated for 16 hrs. in Alexa-555 and Alexa-488-conjugated secondary antibodies (1:100, A-21437 and A-21206, respectively, Invitrogen by Thermo Fisher Scientific; RRID:AB_2535858 and AB_2535792, respectively). Finally, sections were counterstained with Hoechst (1: 200; Millipore, CAS# 875756-97-1).

### Image Acquisition and Analysis

For each UOA-insertion region, we selected sections that contained the largest number of shanks and full shank lengths. These sections were double immunostained for GFAP and NeuN or GFAP and Iba1, and counter stained with Hoechst. High-resolution fluorescent images of these immunostained sections were acquired using an Axio Observer Z1 inverted microscope (Carl Zeiss, Germany) using a 10x objective. Individual images were stitched together using the Zen I software (Carl Zeiss) to obtain a complete wide-field image of the UOA insertion region. For each UOA insertion region, we analyzed one full section per channel (3 channels, 1 for each IHC marker), specifically the section containing the largest number of shanks, using ImageJ-FIJI 1.53f51 software (National Institute of Health, https://imagej.nih.gov/ij/34). Images were transformed into binary black and white images by a thresholding algorithm in ImageJ (https://imagej.nih.gov/ij/docs/guide/146-28.html#toc-Subsection-28.2), and the threshold was then slightly adjusted manually, to closely match the original IHC image. The thresholding separates the pixels within the region of interest (ROI) into those containing signal (which were set to an intensity value of 255, i.e. white) and those belonging to the background (which were set to a value of 0, i.e. black). On these binary images we then calculated the Integrated Density within a 200 x 200 µm window positioned at 4 different cortical depths (corresponding to the cortical layers in **Figs. 4A,6A,8A**) along one edge of each of the tracks left in the tissue by the probe’s shanks. The window size effectively covered the full width of the inter-shank tissue. Integrated Density was defined as the product of the number of white pixels in the measuring window and the maximum pixel intensity value (255), a unit that is referred to as Relative Fluorescence Units (RFUs). This approach effectively measures the number of white pixels in the measuring window. As control, in each section we similarly measured Integrated Density in non-implanted tissue located at least 500µm away from the UOA insertion region (see **Figs. 3,5,7**). These control measurements were used for normalization of the Integrated Density measures shown in **Figures 4,6,8**.

### Statistics

Statistical analyses were conducted using GraphPad Prism 8.0.1 software (San Diego, CA). The datasets were assessed for normality using the Shapiro–Wilk test (for n<50). Parametric tests, including the t-test, Pearson’s correlation, and one-way ANOVA, were utilized when data exhibited a normal distribution. In cases where the normality assumption was violated, non-parametric tests, including the Mann–Whitney test, Kruskal-Wallis test, and/or Spearman correlation, were employed. Data is presented as mean ± standard error of the mean (s.e.m.), and p values <0.05 were considered statistically significant.

## Supporting Information

Supporting Information is available from the Wiley Online Library.

## Acknowledgments

1. E. Fernandez, S. Blair and A. Angelucci contributed equally to this work and share senior authorship. We thank Dr Justin Balsor for help with some experiments. This research was supported primarily by a BRAIN Initiative grant (U01 NS099702) from the National Institute of Health (NIH) to S.B. and A.A. Additional support was provided by grants from the NIH (R01 EY026812, R01 EY031959), the National Science Foundation (IOS 1755431), and the Mary Boesche endowed Chair, to A.A; by an unrestricted grant from Research to Prevent Blindness, Inc. and a core grant from the NIH (P30 EY014800) to the Department of Ophthalmology, University of Utah; by grants PDC2022-133952-100 and PID2022-141606OB-I00 from the Spanish “Ministerio de Ciencia, Innovación y Universidades”, grant CIPROM/2023/25 from the Generalitat Valenciana, and by the European Union’s Horizon 2020 Research and Innovation Programme under Grant Agreement No. 899287 (NeuraViPeR), to E.F.

## Conflict of interest

The authors declare no conflict of interest.

## Data Availability statement

The data that support the finding of this study are available from the corresponding authors upon reasonable request.

## Notes

### Competing Interest Statement

The authors have declared no competing interest.

