## Supplemental Figs 1-5 for "Cortical Response to Acute Implantation of the Utah Optrode Array in Macaque Cortex"

### SUPPORTING INFORMATION

### SUPPORTING FIGURE LEGENDS

**Supporting Figure 1. Statistical comparisons of GFAP immunoreactivity across cortical depths/layers in response to UOA insertion.**

**(A)** Effect of insertion. For each UOA shank diameter group (Left, Middle and Right bar graphs) the raw Integrated Density of GFAP expression (expressed as Relative Fluorescent Units, RFUs, see Experimental Methods) for smooth/round (*s*) and rough/sharp (*r*) UOAs in different layers is statistically compared to each respective control (*ctrl*), using the Mann-Whitney test or t-test, as appropriate. *, **, ***, **** indicate statistical significance at the <0.05, <0.01, <0.001 and <0.0001 level, respectively. **(B)** Effect of shank diameter**.** For smooth/round and rough/sharp UOAs (Left and Right bar graphs) Integrated Density of GFAP expression is statistically compared across UOAs of different shank diameter in different layers, using the Kruskal-Wallis test or one-way ANOVA, as appropriate, corrected for multiple comparisons. Other conventions are as in (A). **(C)** Effect of geometry. For each UOA shank diameter group (Left, Middle and Right bar graphs) Integrated Density of GFAP expression is statistically compared across the two different geometries, smooth/round (*s*) and rough/sharp (*r*) UOAs in different layers, using the Mann-Whitney test or t-test, as appropriate.

**Supporting Figure 2. Double-immunostaining for GFAP and Iba1 in the same sections.**

**(A)** Micrographs of two coronal sections double immunostained for both GFAP and Iba1 at the site of implantation of a 60µm shank diameter smooth/round UOA (Left) and a 60µm rough/sharp UOA (Right). Conventions are as in **Fig. 3A**. Left and right panels show the same sections as in **Fig. 5A** and **5B**, respectively, but here shown in both channels. **(B)** Same as in (A) but for two 85µm shank diameter UOAs. Left and right panels show the same sections as in **Figs. 3C,5C**, and **Figs. 3D,5D**, respectively, but here shown in both channels simultaneously. **(C)** Same as in (A) but for two 100µm shank diameter UOAs. Left and right panels show the same sections as in **Figs. 3E,5E**, and **Fig. 5F**, respectively, but here but here shown in both channels simultaneously. Scale bar: 500µm and valid for all panels

**Supporting Figure 3. Statistical comparisons of Iba1 immunoreactivity across cortical depths/layers in response to UOA insertion.**

**(A)** Effect of insertion. For each UOA shank diameter group (Left, Middle and Right bar graphs) the raw Integrated Density of Iba1 expression for smooth/round (*s*) and rough/sharp (*r*) UOAs in different layers is statistically compared to each respective control (*ctrl*), using the Mann-Whitney test or t-test, as appropriate. *, **, ***, **** indicate statistical significance at the <0.05, <0.01, <0.001 and <0.0001 level, respectively. **(B)** Effect of shank diameter**.** For smooth/round and rough/sharp UOAs (Left and Right bar graphs) Integrated Density of Iba1 expression is statistically compared across UOAs of different shank diameter in different layers, using the Kruskal-Wallis test or one-way ANOVA, as appropriate, corrected for multiple comparisons. Other conventions are as in (A). **(C)** Effect of geometry. For each UOA shank diameter group (Left, Middle and Right bar graphs) Integrated Density of Iba1 expression is statistically compared across the two different geometries, smooth/round (*s*) and rough/sharp (*r*) UOAs in different layers, using the Mann-Whitney test or t-test, as appropriate.

**Supporting Figure 4. Double-immunostaining for GFAP and NeuN in the same sections.**

**(A)** Micrographs of two coronal sections double immunostained for both GFAP and NeuN at the site of implantation of a 60μm shank diameter smooth/round UOA (Left) and a 60μm rough/sharp UOA (Right). Conventions are as in **Fig. 3A**. Left and right panels show the same sections as in **Figs. 3A,7A** and **3B,7B**, respectively, but here shown in both channels. **(B)** Same as in (A) but for two 85μm shank diameter UOAs. Left and right panels show the same sections as in **Fig. 7C** and **7D**, respectively, but here shown in both channels. **(C)** Same as in (A) but for two 100μm shank diameter UOAs. Left and right panels show the same sections as in **Fig. 7E**, and **Figs. 3F**, **7F**, respectively, but here shown in both channels simultaneously. Scale bar: 500μm and valid for all panels.

**Supporting Figure 5. Statistical comparisons of NeuN immunoreactivity across cortical depths/layers in response to UOA insertion.**

**(A)** Effect of insertion. For each UOA shank diameter group (Left, Middle and Right bar graphs) the raw Integrated Density of NeuN expression for smooth/round (*s*) and rough/sharp (*r*) UOAs in different layers is statistically compared to each respective control (*ctrl*), using the Mann-Whitney test or t-test, as appropriate. *, **, ***, **** indicate statistical significance at the <0.05, <0.01, <0.001 and <0.0001 level, respectively. **(B)** Effect of shank diameter**.** For smooth/round and rough/sharp UOAs (Left and Right bar graphs) Integrated Density of NeuN expression is statistically compared across UOAs of different shank diameter in different layers, using the Kruskal-Wallis test or one-way ANOVA, as appropriate, corrected for multiple comparisons. Other conventions are as in (A). **(C)** Effect of geometry. For each UOA shank diameter group (Left, Middle and Right bar graphs) Integrated Density of NeuN expression is statistically compared across the two different geometries, smooth/round (*s*) and rough/sharp (*r*) UOAs in different layers, using the Mann-Whitney test or t-test, as appropriate.

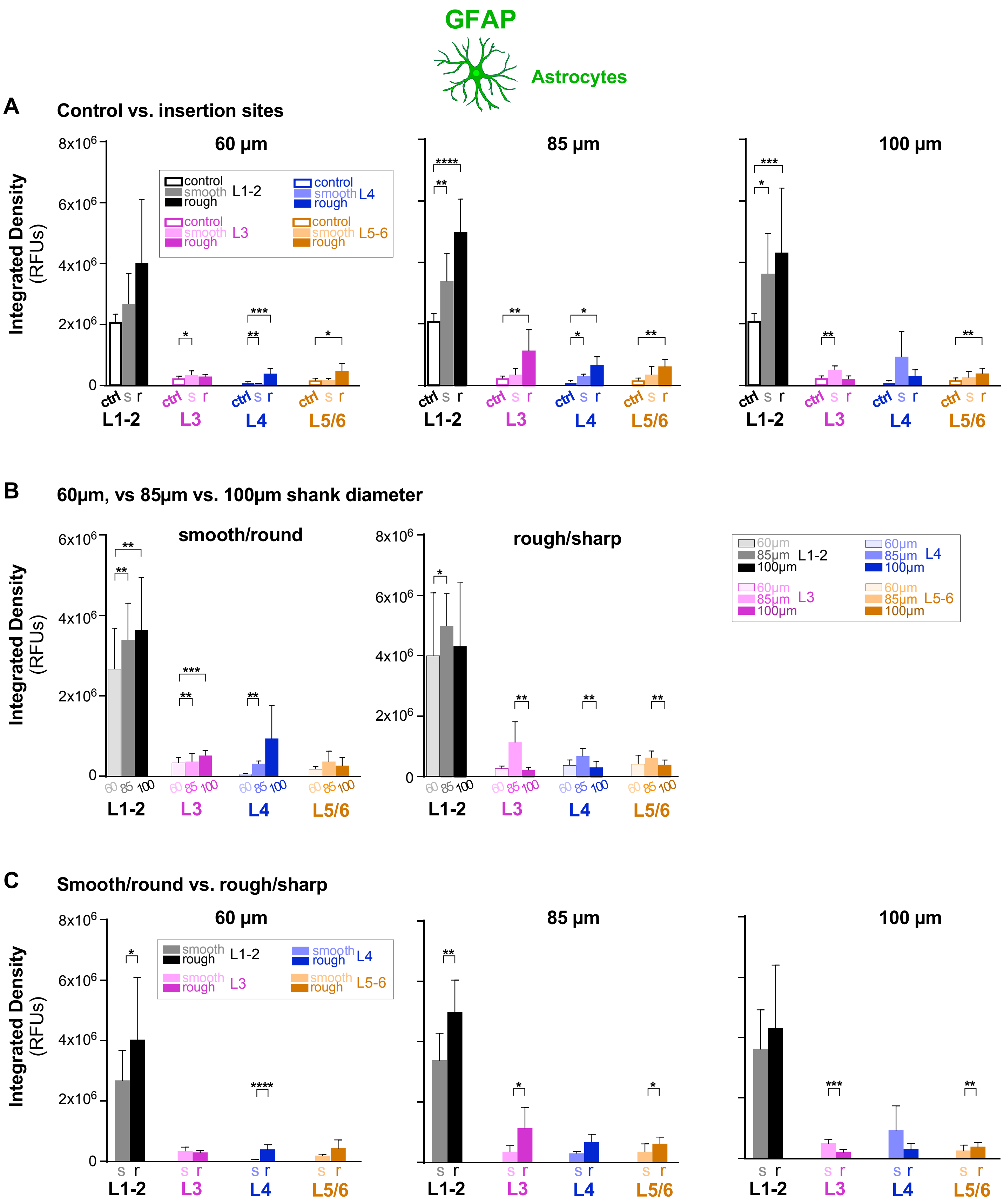


Supporting Figure 1

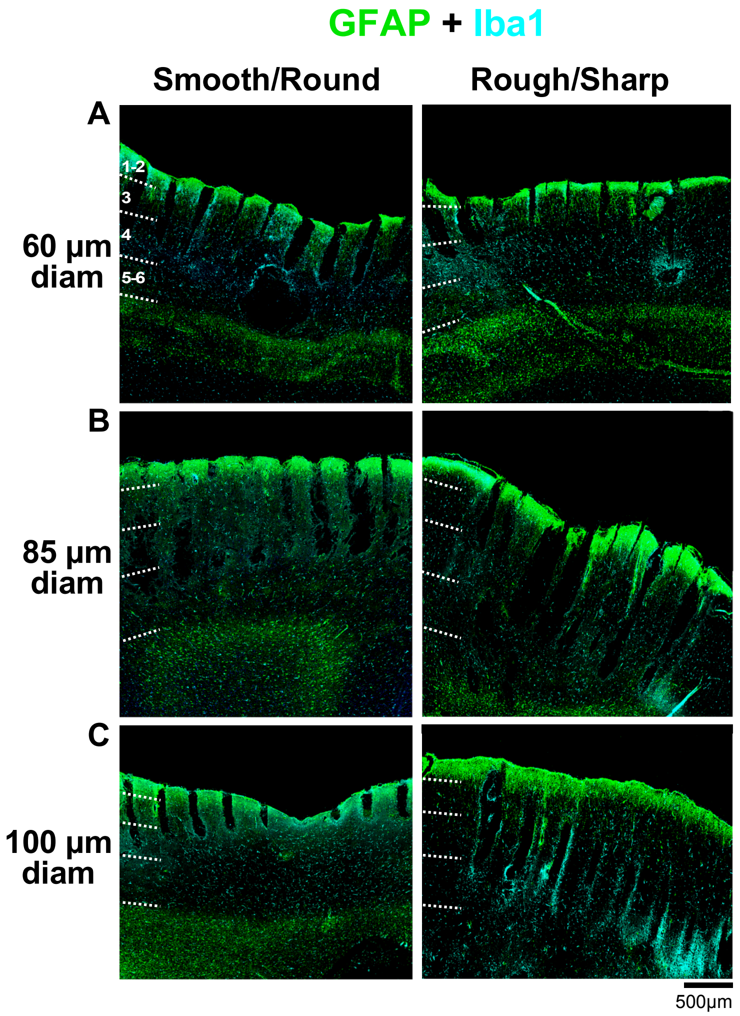


**Suppporting Figure 2**

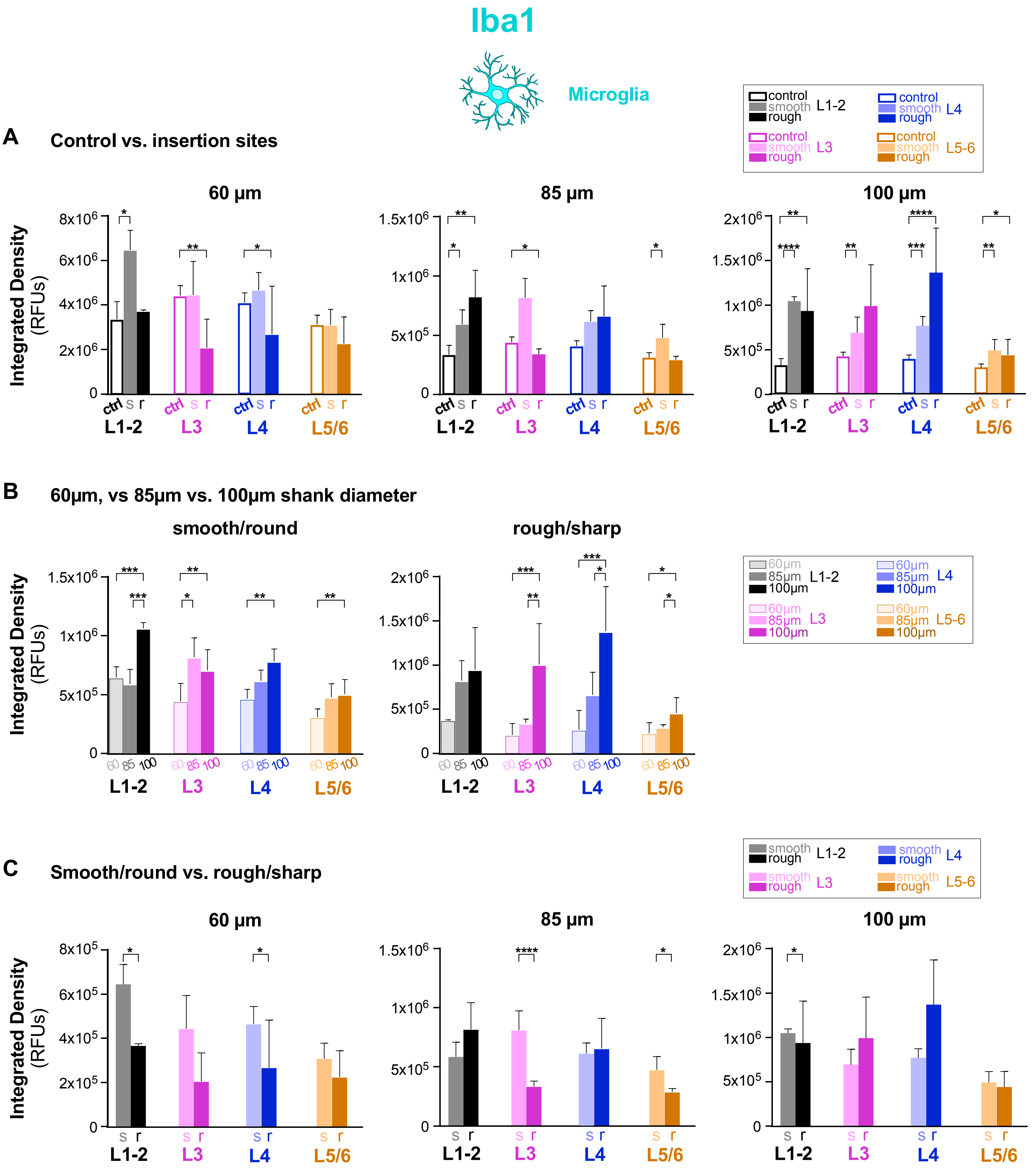


**Supporting Figure 3**

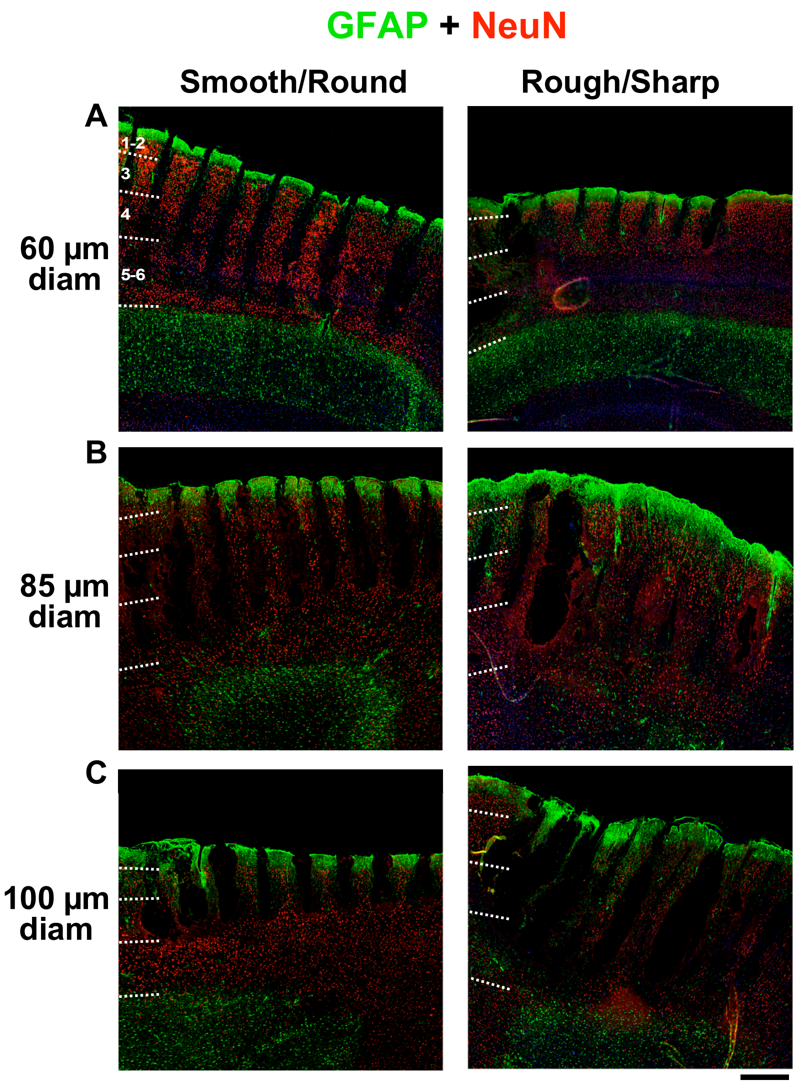


**Supporting Figure 4**

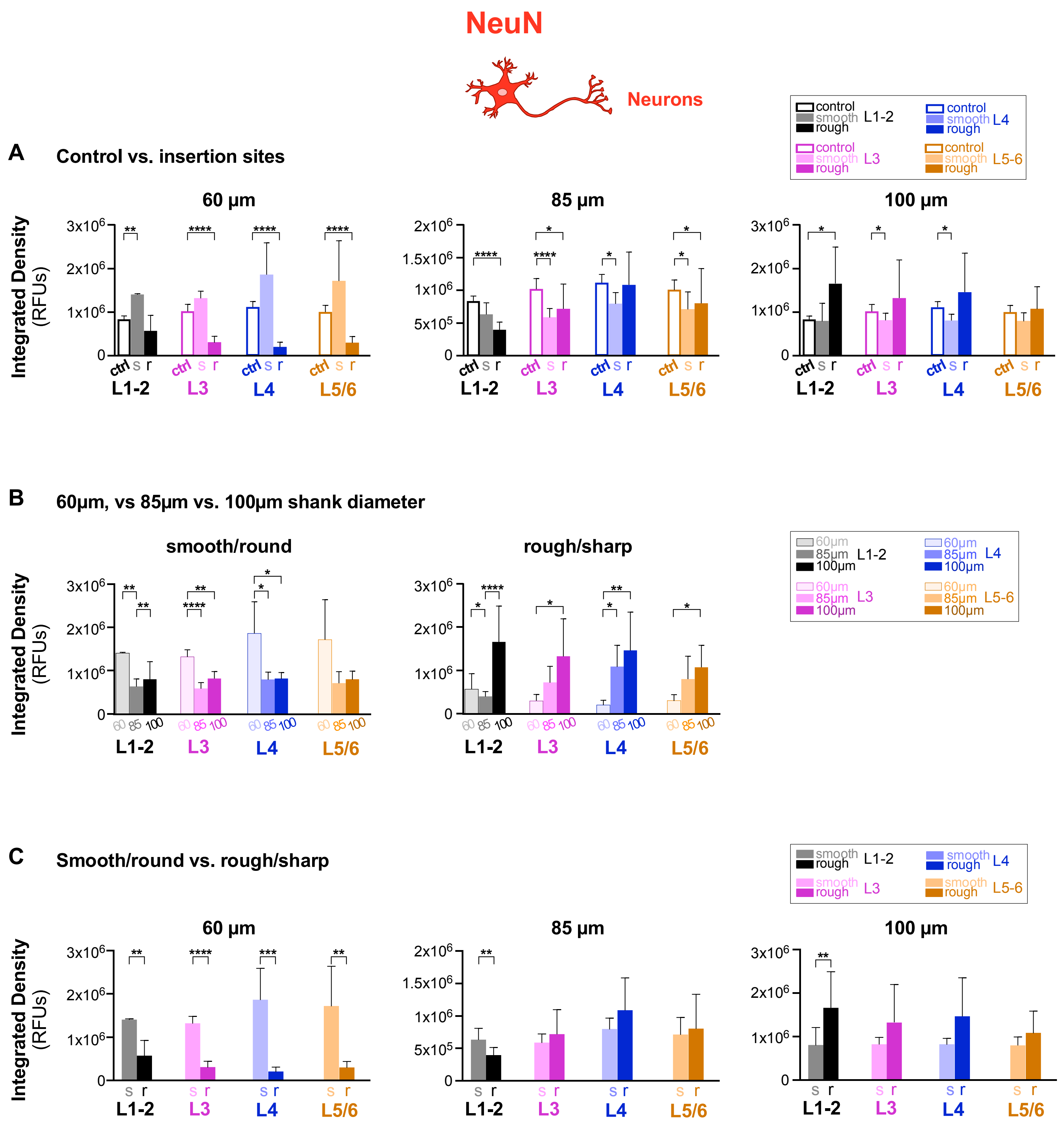


Supporting Figure 5
